# Acetic acid induces osmotic imbalances in drug-resistant bacteria synergistically enhancing cobalt-doped carbon quantum dots bactericidal efficiency

**DOI:** 10.1101/2024.11.06.622380

**Authors:** Adam Truskewycz, Benedict Choi, Line Pedersen, Jianhua Han, Nils Halberg

**Affiliations:** Department of Biomedicine, University of Bergen, Bergen, Norway; Department of Biochemistry and Biophysics, University of California, San Francisco, USA; Department of Biochemistry and Biophysics, University of California, San Francisco, San Francisco, CA, USA; Department of Urology, University of California, San Francisco, San Francisco, CA, USA; Helen Diller Family Comprehensive Cancer Center, University of California, San Francisco, San Francisco, CA, USA; Bakar Computational Health Sciences Institute, University of California, San Francisco, San Francisco, CA, USA; 7Cancer Research Program, QIMR Berghofer Medical Research Institute, Brisbane, Australia

## Abstract

When pathogenic bacteria colonise a wound, they can create an alkaline ecological niche which selects for their survival by creating an inflammatory environment which restricts healthy wound healing to proceed. To aid healing, wound acidification has been exploited to disrupt this process and stimulate fibroblast growth, increase wound oxygen concentrations, minimise proteolytic activity and re-stimulate the host immune system. Within this study, we have developed unique cobalt doped carbon quantum dot nanoparticles which work together with mild acetic acid creating a potent synergistic antimicrobial therapy. The acidic environment alters the osmotic balance of microorganisms forcing them to swell and speed up the internalisation of the ultra-small particles. The particles hyperpolarise the bacterial membranes and generate damaging peroxidase species resulting in cellular lysis. In mice, cobalt doped carbon quantum dots remove MRSA infection while allowing wounds to heal at equivalent rates to uninfected wounds. This work demonstrates how synergistic antimicrobial treatment strategies can be successfully used to combat antimicrobial resistant infections.

## Main text

There are currently > 4.5 million deaths associated with bacterial antimicrobial resistance (AMR) globally arising from either a direct infectious disease, or through additive physiological stress experienced by those already suffering from pre-existing conditions (i.e. cancer, organ transplantation, HIV, liver and kidney disease, diabetes etc.). Antimicrobial resistance is an increasingly serious threat with too few drugs in the developmental pipeline (WHO)^1^ adept to combat the most adaptive and resilient species. Most new drug candidates are variations of pre-existing commercial chemical antibiotics; however, there is pressing need for innovative treatments. There are currently only 2 drugs in the pipeline fulfilling at least one of the WHOs four innovation criteria which includes: drugs with a new chemical class, new microbial targets, new mechanism of action, and/or free from cross antimicrobial resistance^2^

Ultrasmall nanoparticles hold great promise as antimicrobial agents having the potential to address several of the WHO antimicrobial innovation criteria as they possess properties of both molecular/chemical and physical states ^3^. Their structure separates them from traditional chemical drugs and provides a platform for multifunctional activity which can further challenge resistance. Carbon quantum dot nanoparticles are a class of ultrasmall nanoparticles. Unlike other rigid carbon nanoparticles (carbon nanotubes, graphene etc.), they possess high hydrophilicity and monodispersity making them less likely to be bioaccumulated and have shown a high degree of biocompatibility in numerous studies^4–6^. Their carbon core contains varying organic chemical structures providing them with abundant functionalization opportunities (i.e. metal dopants and/or organic ligands), to instill them with multiple functional characteristics^7^. Despite their potential, their ability to remove infection *in vivo* and restore pathogen infected wounds remains unexplored.

Healthy wound healing proceeds through four main phases namely hemostasis, inflammation, proliferation, and remodeling. During these stages, the environments pH fluctuates with i) an acidic inflammation stage which reduces microbial colonization and promotes vascular regeneration, ii) a marginally alkaline granulation stage (pH 7.0 – 7.5) to promote cell proliferation and skin remodeling, followed by a iii) re-acidification of the environment to restore the skins acid mantle^8^. However, infected wounds often remain fixed in the inflammatory state through the action of persistent pathogenic microorganisms. Preventing these microbial infections from forming is key to ensuring healthy wound healing processes^9^.

Pathogenic microorganisms create an environmental niche which is beneficial for their survival whilst inhibiting the host’s immune system from functioning to control their presence^10^. An alkaline pH (pH 7.0 – 9.0)^11^, reduced wound oxygen content,^12^ and increased temperature are frequently associated with seriously infected wounds^13^. Healing wounds undergo natural acidification, producing lactic acid, which minimises wound infection, supports vascular migration, DNA replication, wound oxygenation, collagen formation immune activity and growth of blood vessels^14^. Therefore, wound acidification has been suggested as a viable approach for reducing infection^15^. Acetic acid is considered a mild acid, and at diluted concentrations, has been shown to have pro-wound healing properties^16^. It has also been used against the plague^17^ and has in recent days, been studied for its antimicrobial activity and it is particularly effective against pathogenic *Pseudomonas aeruginosa*^18^, however it shows negligible bactericidal activity against many other wound colonising bacterial species at concentrations ^19^ below the threshold for dermal discomfort (3%). Its short residence time in the wound environment (∼1 h)^20^ allows it to temporarily adjust the wound environment without bioaccumulating, allowing healthy wound healing associated pH fluctuations to continue.

Here we have developed a combination therapy utilising weak acetic acid (0.06%, pH 5.5) together with novel, ultra-small, cobalt doped carbon quantum dot nanoparticles (Co-CQD) as a potent antimicrobial strategy against several pathogenic species including methicillin and oxacillin resistant *Staphylococcus aureus* (MRSA), *Escherichia coli* **(Figure 1**), and *Enterococcus faecalis*. A weak acetic acid environment creates osmotic disruption resulting in cell swelling and more rapid Co-CQD particle uptake. The particles then destroy bacteria cells through the generation of peroxides and hyperpolarises their membranes. The particles show a high degree of biocompatibility with dermal fibroblasts at concentrations well over the minimum bactericidal concentrations and were able to remove multi drug resistant MRSA infections from mouse wounds without reducing wound healing rates. Together, the platform highlights a biocompatible and efficient and strategy for combatting bacterial infections *in vitro* and *in vivo* by blending traditional and innovative combination strategies.

**Figure 1:**
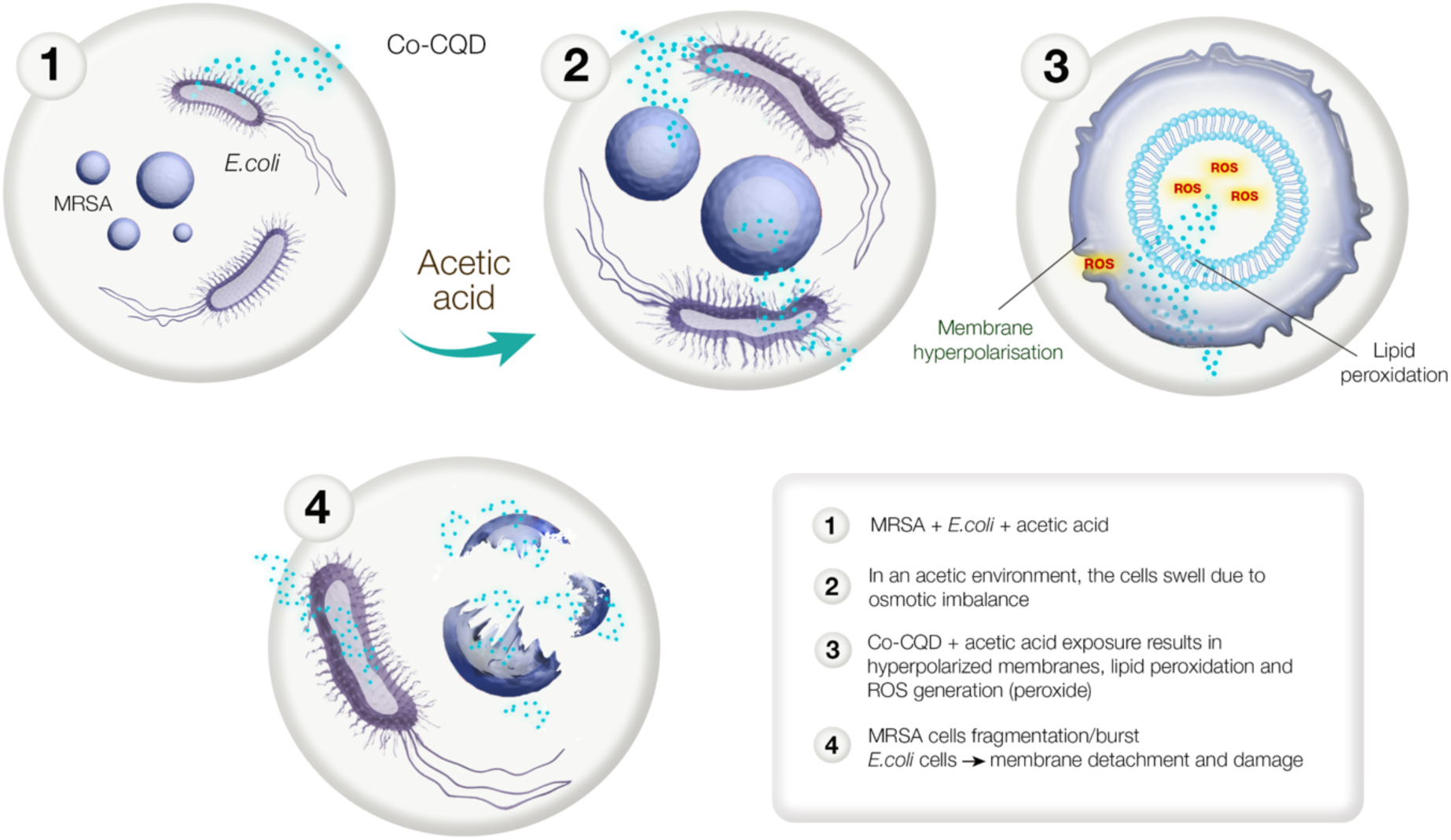
Schematic overview highlighting how the addition of dilute acetic acid (pH 5.5) results in microbial cell swelling and increases the rate of Co-CQD nanoparticle uptake. The Co-CQD particles in this environment hyperpolarize bacterial cell membranes, damage cellular lipids, and create ROS (peroxides) which collectively destroy the bacterial cell.

### Synthesis and characterisation of Co-CQDs

Carbon quantum dots doped with cobalt were designed as potent antimicrobial nanoparticles due to cobalt’s documented antimicrobial activity^21^ along with carbon quantum dots biocompatibility and potential for functional chemical modifications^22–24^. Transmission electron microscopy (TEM) of the particles demonstrated that they were monodispersed, and had sizes ranging between 1.2 – 4.0 nm, with an average size of 2.6 nm (**Figure 2A, B**). The hydrothermal conditions during CQD synthesis facilitated the integration of hexamine cobalt chloride (HACC) derived nitrogen and cobalt into the carbon quantum dots core matrix. XPS analysis showed the particles to be comprised of C (64.7 At%), Co (2.3 At%), N (3.3 At%), O (27.5 At%), Cl (0.3 At%), and other (1.9 At%: **Figure 2C**).

**Figure 2:**
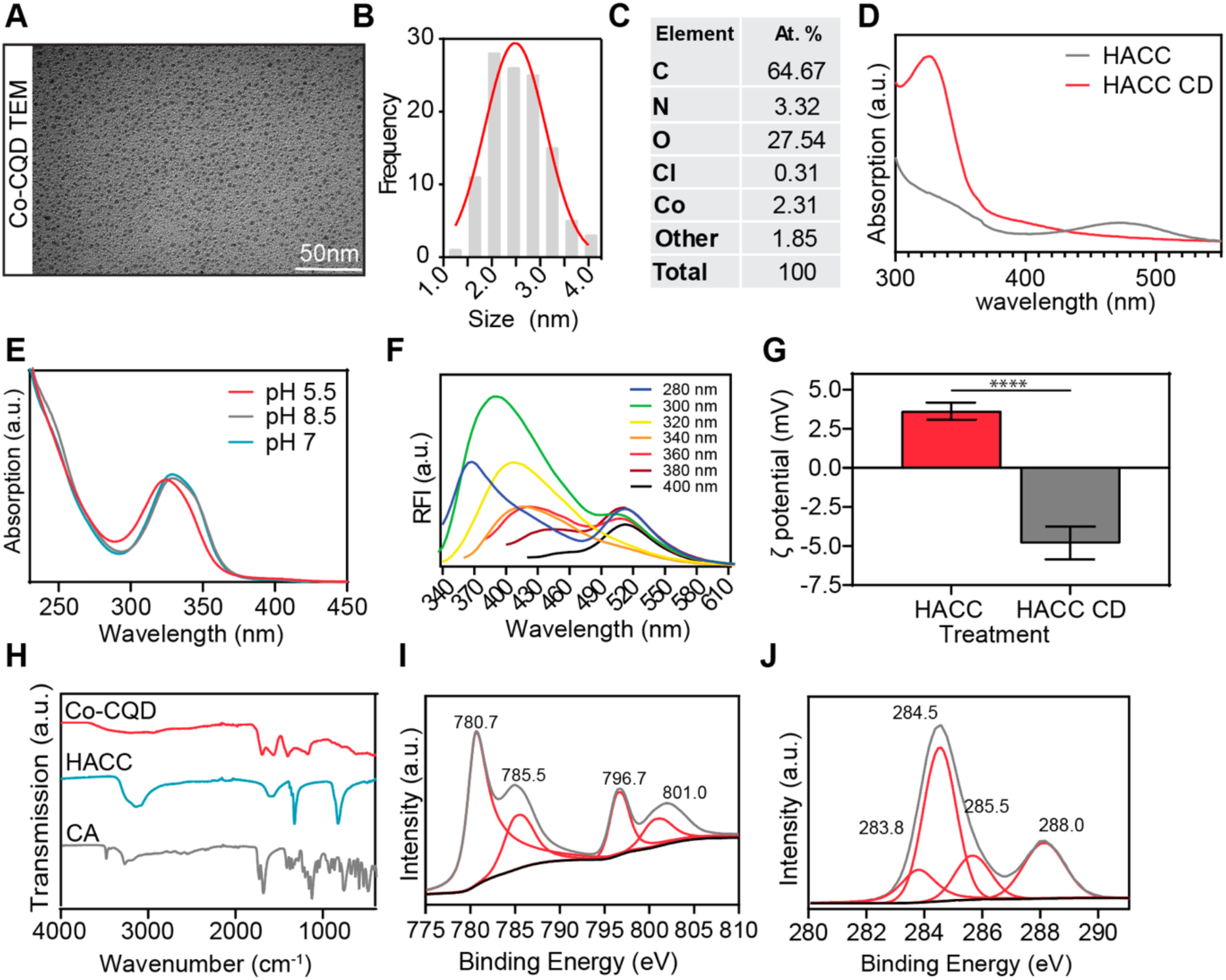
Co-CQD characterisation. **a**, Representative TEM images of Co-CQD nanoparticles. Scale bar is 50 nm. **b**, Histogram of Co-CQD sizes produced through hydrothermal synthesis conditions. **c**, XPS elemental analysis of Co-CQDs (At%). **d**, Absorbance spectra of Co-CQD and its synthesis precursor hexamine cobalt chloride solution. **e**, Absorbance of Co-CQDs in acidic, basic and alkaline conditions. **f**, Excitation and emission fluorescence spectra of Co-CQDs. **g**, Zeta potential of hexamine cobalt chloride solution and Co-CQD solution. **h**, FTIR spectra of Co-CQDs and the precursors used to synthesize it, namely citric acid and hexamine cobalt chloride. i, X-ray photoelectron spectroscopy (XPS) Co2p peaks from Co-CQDs. **j**, XPS C1s peaks from Co-CQDs. Error bars represent the standard deviation. Statistical significance was determined using (e) Unpaired t test (two tailed). **** signifies p < 0.0001

The absence of the parental HACC compound on the CQDs is observed by differences in their absorbance spectra with the particles possessing a strong peak at 330 nm whilst losing HACC characteristic absorbance peak at 435 nm (**Figure 2D**). The particles show slight blueshift in absorbance in acidic conditions indicating breaks in structural conjugation (**Figure 2E**). The fluorescent particles possess a maximum fluorescence emission at λ_max_ 390 nm when excited at 300 nm light and incremental red shifting of the peak emissions occurred with increasing excitation wavelengths between 280 – 400 nm (**Figure 2F**). Variations in zeta potential of HACC solution and Co-CQDs were shown to switch from a slight positive to slightly negative charge (3.6 vs −4.8 mV) indicating carboxylic acid functionalization derived from the citric acid precursor (**Figure 2G**). Nevertheless, these low zeta potential values indicate the particles did not possess strong surface charges. The hexamminecobalt (III) chloride precursor possessed FTIR peaks at 3250, 1615, 1325 and 823 cm^−1^ which are all representative of N-H bonds. However, these functionalities were absent from the nanoparticles. The nanoparticles displayed carboxylic acid functionalities shown by FTIR spectra peaks at 3300-2500 cm^−1^ (O-H stretch), 1690 cm^−1^ (C=O stretch), 1400 cm^−1^ (O-H bend), and 1227 cm^−1^ (C-O stretch) (**Figure 2H**).

The Co-CQDs possess a broad Raman D band between 1315-1375 cm^−1^ representing a disordered sp^2^ carbonaceous material and a broad Raman G band between 1510 – 1545 cm^−1^ attributable to amorphous carbon (**Supplementary Figure 2A**). The Co 2p XPS spectrum shows two dominant Co^2+^ peaks centred at 780.7 and 796.7 eV for 2p3/2 and 2p1/2 respectively. Corresponding satellite peaks are centred at 785.5 and 801.0 respectively (**Figure 2I**). These binding energies coupled with that of the O1s at 531.1 are characteristic of Co(OH)_2_ (**Supplementary Figure 2B**). The particles possess nitrogen containing ring structure as indicated by XPS peaks at 399.1 and 399.8 eV representative of lactam or cyano nitrogen, and pyrrolic nitrogen respectively (**Supplementary Figure 2C**). Carbons C1s peaks at binding energies of 283.8, 284.5, 285.5, 288.0 eV (**Figure 2J**) are assigned to sp2 and sp3 hybridized carbon, C-OH, and C=O respectively.

These characterisations reveal that the fluorescent Co-CQD are monodisperse ultra-small carbonaceous particles doped with nitrogen and cobalt heteroatoms. The particles core is disordered, and its surface contains carboxylic acid functionalities. The parental synthesis precursor (HACC) is absent from the final nanoparticle product with hydrothermal conditions transforming Co^3+^ into a Co^2+^ oxide.

### Acetic acid Co-CQDs exhibit synergistic antimicrobial activity

Methicillin and oxacillin resistant *Staphylococcus aureus* (MRSA, ATCC 43300), *Escherichia coli* (ATCC 25922), and vancomycin-sensitive *E. Faecalis* (ATCC 29212) are amongst the most frequently identified microorganisms in infected wounds^25–27^ and were therefore chosen to investigate the antimicrobial activity of Co-CQD nanoparticles. Acetic acid has been assessed for its antimicrobial activity in pre-clinical trials and has shown strong antimicrobial activity against the gram-negative Pseudomonas aeruginosa^28, 29^. It has anti-biofilms forming properties and its undissociated form possesses a neutral charge allowing it to freely cross bacterial membranes causing cellular acidification and osmotic imbalances. It has been shown to slow the growth of pathogenic bacteria, but its bactericidal activity against these strains is limited. Due to its low dermal toxicity (LD_50_ of 1.06 g/kg in rats), cellular internalisation properties and bacteriostatic nature, acetic acid was selected as a medium for synergistic Co-CQD antimicrobial activity.

The Co-CQD nanoparticles exhibited strong antimicrobial activity towards all strains in aqueous growth cultures, particularly in a weak acetic acid environment (0.06%) (**Figure 3A**). Acetic acid concentrations of 1-2% (pH ∼2.4) have been well tolerated in preclinical trials^30^, however in this study, we have kept the pH at levels naturally found in healthy healing wounds^31^. At pH 5.5, all MRSA, *E. coli* and *E. faecalis* cells were destroyed at a 38, 75, and 75 µg/ml Co-CQD concentration respectively (minimal bactericidal concentration: MBC) (**Figure 3A, Supplementary Figure 3B**). The MIC determined by OD_600_ of antimicrobial overnight culture was 38, 38 and 9 µg/ml respectively indicating a bacteriostatic influence on *E. coli* and *E. faecalis*. At pH 7.0, a strong but reduced antimicrobial activity of Co-CQD was observed indicating the particles exploit the acidic environment for enhanced antimicrobial activity (**Figure 3A**). At pH 7.0 MRSA, *E. coli* and *E. faecalis* possessed an MBC of 150, 600 and 300 µg/ml respectively and an MIC of 75, 300 and 150 µg/ml respectively (**Figure 3A**). An alkaline environment of pH 8.5 further decreased the antimicrobial activity of the particles (**Supplementary Figure 3A**).

**Figure 3:**
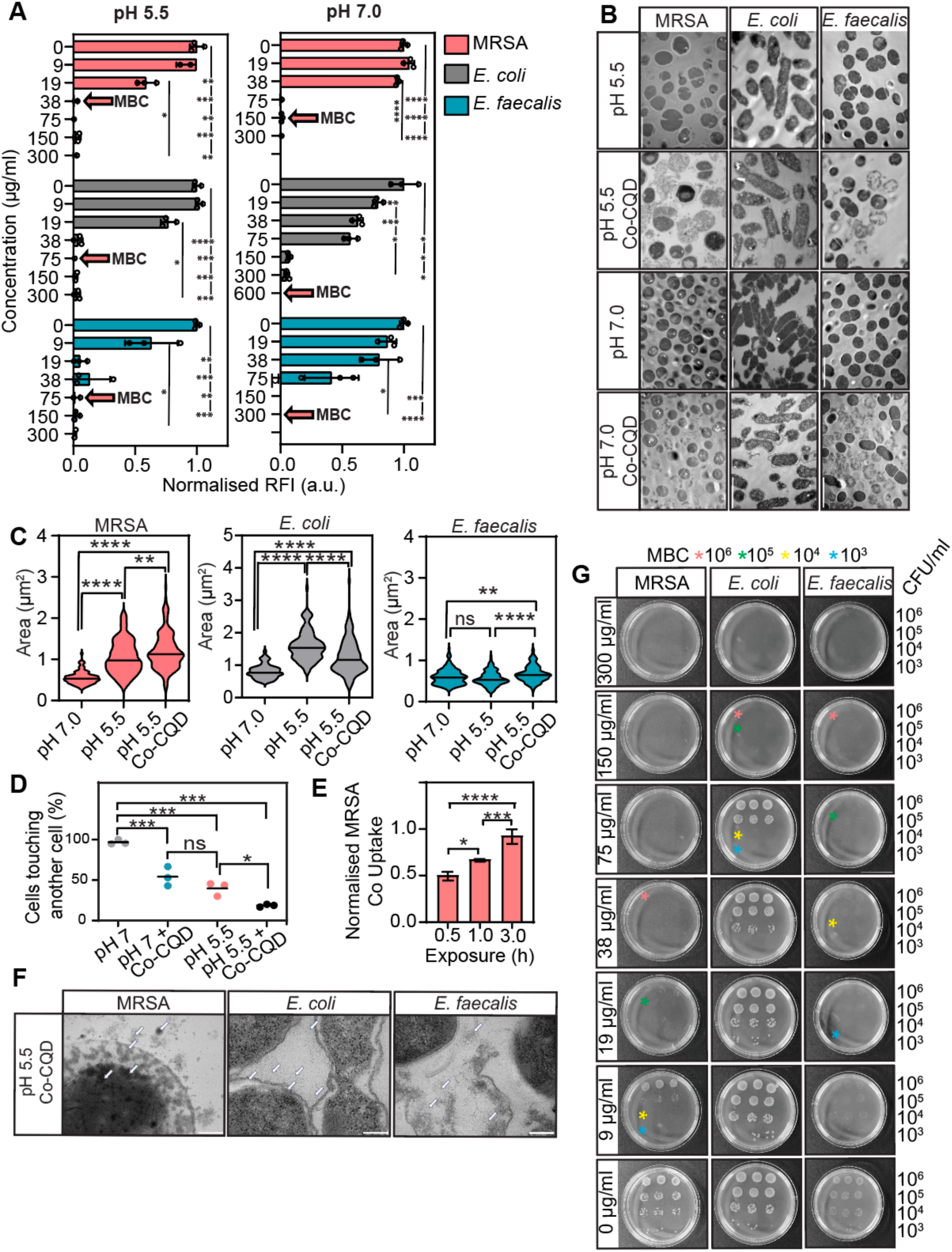
Antibacterial activity of Co-CQDs. **a**, Optical density (OD_600 nm_) of MRSA, *E. coli*, and *E. faecalis* in pH adjusted liquid growth cultures after 24 h exposure to Co-CQDs. Red arrows denote the MBC Co-CQD concentration. **b**, TEM images of MRSA, *E. coli*, and *E. faecalis* at pH 5.5, pH 5.5 + Co-CQDs, pH 7.0, and pH 7.0 + Co-CQDs following a 24 h exposure period. **c**, MRSA, *E. coli*, and *E. faecalis* cell sizes when exposed to nutrient broth pH 7.0, nutrient broth + acetic acid (pH 5.5) and nutrient broth + acetic acid (pH 5.5) + Co-CQDs. **d**, Ratio of MRSA peptidoglycan content relative to cell size. **e**, TEM of Co-CQD interaction with MRSA, *E. coli* and E. faecalis cell membranes at pH 7 and 5.5. **f**, Number of *E. coli* cells that are touching another *E. coli* cell (cell-cell interaction) at pH 7 and 5.5, with and without Co-CQD treatment. **g**, Growth of MRSA, *E. coli*, and *E. faecalis* on nutrient agar plates supplemented with differing Co-CQD concentrations (0 – 300 µg/ml) at pH 5.5 (acetic acid adjusted). Each plate has been seeded with different concentrations of bacteria (5.0 × 10^6^, 10^5^, 10^4^, and 10^3^ CFU/ ml) from top to bottom in triplicate. Stars of red, green, yellow, and blue represent the MBC for 10^6^, 10^5^, 10^4^, and 10^3^ CFU/ ml respectively. Error bars represent the standard deviation. Statistical significance was determined using (a) a two-way analysis of variance (ANOVA) with Tukey’s multiple comparison test or a (d) one-way ANOVA with Tukey’s multiple comparison test. ns, *, **, ***, **** signifies not significant, p < 0.05, p < 0.005, p < 0.0005 and p < 0.0001, respectively.

To better understand the synergistic antimicrobial function between Co-CQDs and acetic acid, we first assessed the effect of Co-CQD on bacterial morphology by transmission electron microscopy under various pHs. The addition of Co-CQDs to MRSA and *E. faecalis* cells in the acetic acid environment resulted in detachment of the cytoplasmic membrane from the outer membrane, release of cytoplasm constituents and complete cellular disintegration (**Figure 3B**). *Escherichia coli* cells exposed to Co-CQDs (in neutral and acidic conditions) had obvious detachment of the cell wall from their cytoplasm (**Figure 3B**). At this ultrastructural level, both MRSA and *E. coli* swelled to nearly double the size when grown at pH 5.5 compared to pH 7.0 (1.91 and 1.99 times respectively) indicating osmotic imbalance in these cells (**Figure 3C**). In MRSA, there was no significant difference between the ratio of cell size to peptidoglycan thickness between pH 7 and 5.5, however, as the cells were twice the size in acetic acid conditions, there was twice the amount of peptidoglycan per cell (**Supplementary Figure 3D**). Acidic acid conditions also prevented cell-cell interactions in *E. coli* with the percentage of cells touching a neighbouring cell dropping from 96.7 % at pH 7 to 39.7 % at pH 5.5. The addition of Co-CQDs further reduced cell-cell interactions with 54.2 % at pH 7 and 18.5% at pH 5.5 (**Figure 3D**). An increase in Co-CQD uptake by MRSA cells was confirmed through cellular cobalt uptake. At both pH 5.5 and 7.0, particles were able to translocate the membrane with particle uptake being more rapid at pH 5.5 (**Figure 3E**). TEM images focused on the cell membrane at 60,000 x magnification show that at pH 5.5, Co-CQDs translocate through the MRSA cell membrane and into the cell (white arrows). In these conditions, *E. faecalis* cells showed Co-CQD internalisation resulting in cell lysis (**Figure 3F**). The gram-negative *E. coli* cell membrane seems largely impermeable to nanoparticle translocation; however, damaged membranes brought upon from the particles and the acetic acid environment presented a route for monodispersed Co-CQD entry into the cell (white arrows).

To understand how Co-CQDs influence bacterial colonisation on solid surfaces, the nanoparticles were incorporated into agar plates at pH 5.5 and 7.0. As seen with the liquid cultures, an acetic acid environment provided enhanced antimicrobial activity (**Figure 3G**). The MBC for MRSA in acidic conditions was dependent on initial bacterial seeding concentration with no growth present at Co-CQD concentrations of 38, 19 and 9 µg/ml for 5.1 × 10^6^, 10^5^ and 10^4^ CFU/ml respectively. In neutral conditions, this rose to 150 µg/ml regardless of bacterial seeding concentration (**Supplementary Figure 3C**). The MBC for *E. coli* in acidic conditions was 150, 150 and 75 µg/ml for 6.6 X 10^6^, 10^5^ and 10^4^ CFU/ml respectively and for *E. faecalis* it was 150, 75 and 38 µg/ml for 5.5 X 10^6^, 10^5^ and 10^4^ CFU/ml respectively (**Figure 3G**). Additionally, *E. faecalis* exhibited obvious growth suppression as seen by faded colonies across all treatment concentrations.

These experiments thus reinforce potent antimicrobial efficiency of Co-CQD across both gram positive and negative bacteria, in solid and aqueous settings, particularly in a mild acetic acid environment. Acetic acid was shown to disrupt the osmotic balance of both gram-positive (MRSA) and gram-negative (*E. coli*) cells causing them to swell. In this environment, the action of Co-CQDs is greatly enhanced with gram positive cells actively translocating particles into the cell and gram-negative cells allowing particle translocation through damaged membranes.

Having demonstrated potent antibacterial activity in *in vitro* assays, we next sought to assess their functionality *in vivo*. To investigate this, wounds (5 mm diameter) were created onto the back of female black six mice, infected with MRSA bacteria (1.3 × 10^7^ CFU) and treated with Co-CQD nanoparticles (0.87 mg/kg) (**Figure 4A**). After 24 hours, wounds inoculated with MRSA possessed visibly obvious infection and the presence of 6.2 × 10^8^ CFU/g (**Supplementary Figure 4A, B**). The addition of Co-CQD nanoparticles greatly reduced the bacterial load by 4.3 log (3.0 × 10^4^ CFU/g = 99.995% reduction) indicating potent acute *in vivo* antimicrobial activity. To further assess Co-CQD pathogen control, and to investigate their impact on wound healing, infected wounds were treated with Co-CQD, and wound infection and closure rates were determined over 7 days. The addition of Co-CQDs to MRSA infected wounds (1.09 mg/kg) removed all bacterial infection (**Figure 4B, C**).

**Figure 4:**
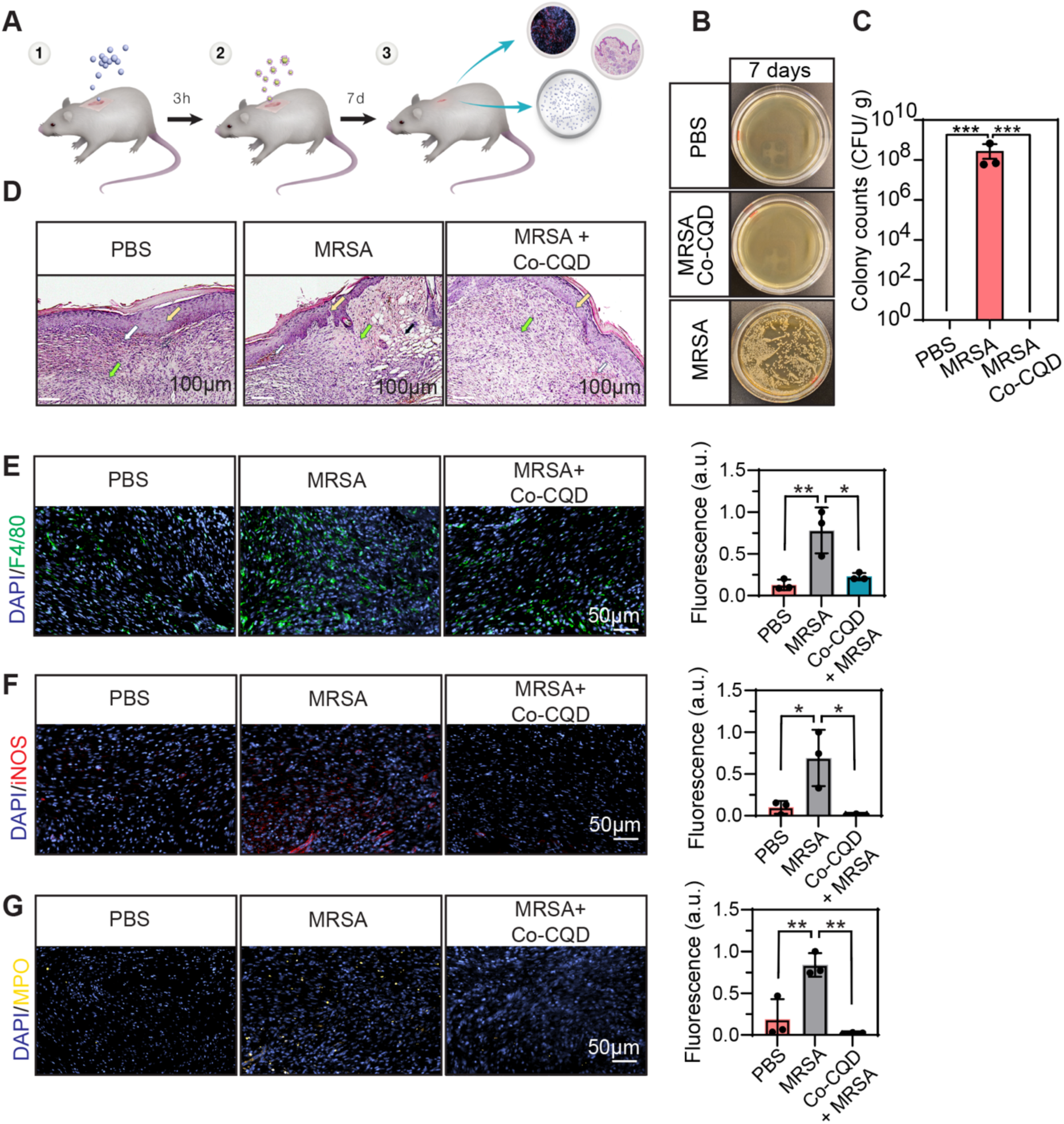
Co-CQD treatment removes infection in vivo, restores skin physiology, and reduces bacterial characteristic immune responses. **a**, Schematic representation of in vivo antimicrobial treatment workflow showing bacterial infection of wounds, addition of Co-CQD nanoparticles, 7 day wound healing period followed by wound microbial load analysis, IF and histological analysis. **b**, agar plates representing wound bacterial load after 7 d treatment exposure. **c**, quantification of bacterial presence in wound areas following treatment after 7 d. **d**, Representative H&E stained wound regions of PBS, MRSA infected, and Co-CQD treated MRSA infected wounds. Green arrows denote granulation tissue, white arrows denote neovascularisation, yellow arrows denote epithelium, and black arrows denote subepidermal haemorrhage tissue. **e**, F4/80 macrophage presence in wound areas following treatment after 7 d. **f**, iNOS inflammatory macrophage presence in wound areas following treatment after 7 d. **g**, MPO neutrophil presence in wound areas following treatment after 7 d. Error bars represent the standard deviation. Statistical significance was determined using (a, b) a two-way analysis of variance (ANOVA) with Tukey’s multiple comparison test or a (e, f) one-way ANOVA with Tukey’s multiple comparison test. ns, *, **, ***, **** signifies not significant, p < 0.05, p < 0.005, p < 0.0005 and p < 0.0001, respectively.

Hematoxylin and eosin stain (H&E) staining showed that the MRSA infected wounds possessed subepidermal haemorrhage tissue areas (black arrows), non-uniform granulation (green arrows), and an incomplete epidermis layer (yellow arrows) despite also possessing clear neovascularisation (white arrows) (**Figure 4D**). This tissue damage suggests that wound inflammation and immune presence is active which may be associated with increased MRSA presence. Treatment with Co-CQD nanoparticles resulted in normalization of the wound areas including abundant granulation tissue (green arrows), structurally complete epidermis layers (yellow arrows) and clear neovascularisation (white arrows) (**Figure 4D**).

Immunofluorescence staining demonstrated that Co-CQDs treatment of MRSA infected wounds contained significantly reduced macrophage (F4/80, **Figure 4E**), inflammatory macrophage iNOS, (**Figure 4F**) and neutrophil (MPO, **Figure 4G**) positive cells indicating decreased inflammation as a result of reduced bacterial infection. As the presence of pathogenic bacteria increases iNOS activity^32–34^, its decreased presence in Co-CQD treated groups supports their antimicrobial activity. There was no significant difference in F4/80, iNOS and MPO positive cells between the Co-CQD treatment and the PBS control.

Combined, these findings demonstrate that the particles removed drug resistant infection from mice wounds resulting in reduced bacterial characteristic immune presence leading to a skin pathology comparable to the uninfected healing wounds.

### Acetic acid primes MRSA for Co-CQD-dependent oxidative damage

Having demonstrated entry into the bacteria, we next set out to determine the mechanisms underpinning the bactericidal effects of Co-CQD treatment. To this end, we next performed RNA sequencing on bacteria exposed to pH 7.0, 5.5, and pH 5.5 with Co-CQDs. The volcano plot in **Figure 5A** highlights that MRSA exposed to acidic conditions responded by upregulation of 84 and suppressed expression of 87 genes. When Co-CQDs were added to these acidic conditions there were further 53 upregulated and 67 downregulated genes. Pathways analysis of MRSA in an acetic acid environment highlighted repressed genes belonging to oxidative phosphorylation indicating that ATP generation was supressed, resulting in reduced cellular respiration (**Figure 5B**). This was supported by a decreased metabolic activity measured by the cells inability to convert non fluorescent resazurin to fluorescent resorufin by dehydrogenase enzymes found in metabolically active cells (**Figure 5C**). Genes relating to hydrogen ion transport were also repressed as an adaptive measure for tolerating the acidic environment.

**Figure 5:**
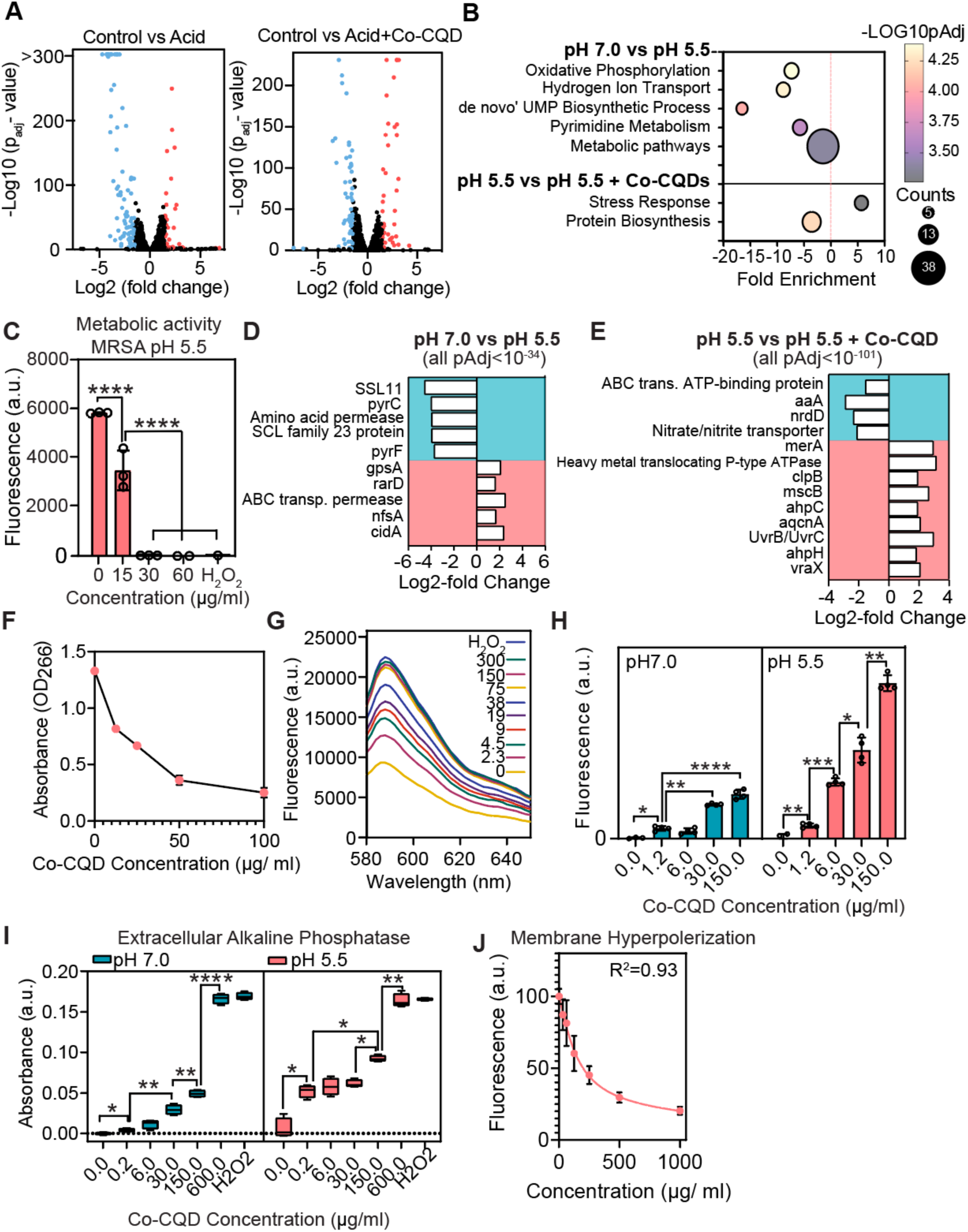
Mechanism of Co-CQDs Antibacterial activity. **a,** Volcano plots of differentially expressed genes between MRSA subjected to pH 7.0 vs pH 5.5 (acetic acid adjusted). and differentially expressed genes between MRSA subjected to pH 5.5 vs pH 5.5 + Co-CQDs. **b**, pathways analysis plot of MRSA subjected to pH 5.5 vs pH 5.5 + Co-CQDs. **c**, Metabolic activity (Resazurin conversion to resorufin) of MRSA when subjected to Co-CQDs at different concentrations at pH 5.5 (acetic acid adjusted). **d**, Expression of significant genes when MRSA is transferred from pH 7.0 to pH 5.5 (acetic acid adjusted). **e**, Expression of significant genes when MRSA is transferred from pH 5.5 (acetic acid adjusted).to pH 5.5 containing Co-CQDs. **f**, ROS generation determined through the oxidation of ascorbic acid at pH 7.0. **g**, Hydrogen peroxide generation of Co-CQDs measured via the Amplex Red assay. **h**, Co-CQDs superoxide generation capacity and its interaction with nucleic acids in a pH 7.0 and 5.5 environment. **I**, alkaline phosphatase activity indicating MRSA membrane damage at pH 7.0 and 5.5 from Co-CQD exposure. **j**, Membrane hyperpolarisation of MRSA at pH 5.5 resulting from differing Co-CQD exposure. Error bars represent the standard deviation. Statistical significance was determined using (a, b) a two-way analysis of variance (ANOVA) with Tukey’s multiple comparison test or a (e, f) one-way ANOVA with Tukey’s multiple comparison test. ns, *, **, ***, **** signifies not significant, p < 0.05, p < 0.005, p < 0.0005 and p < 0.0001, respectively.

The addition of Co-CQD to MRSA in an acetic acid environment increased the regulation of bacterial stress responses and decreased protein biosynthesis pathways (**Figure 5B**). Of the genes identified in the stress response pathway, DnaK, DnaJ and GrpE chaperones, Clp ATPases (clpP and clpB), and CtsR and HrcA regulons been implicated in cellular protection from oxidative stress^35–37^.

At the single gene level, when MRSA was exposed to an acetic acid environment, several genes responsible for drug exporting, antibiotic resistance and chemical detoxification were upregulated (EamA family transporter (RarD), ABC transporter permease, NAD(P)-dependent oxidoreductase, oxygen-insensitive NADPH nitroreductase etc.) indicating that the cells were trying to establish a new favourable equilibrium in the adverse acidic environment (**Figure 5D**).

When Co-CQD nanoparticles were added to acetic acid growth media, enhanced bacterial stress responses were observed. Several genes responsible for the protection against oxidative stress were upregulated (ATP-dependent chaperone (ClpB), protein arginine kinase (mcsB), alkyl hydroperoxide reductase subunit C (ahpC), hypothiocyanous acid reductase (MerA), aconitate hydratase (acnA)) indicating that the particles were generating ROS (**Figure 5E**). In addition, alkyl hydroperoxide reductase subunit F (ahpF), was upregulated which is responsible for protecting DNA from peroxides and C1q-binding complement inhibitor VraX emphasizes that cell wall stress was apparent due to Co-CQD particles. Several heavy metal translocating genes were upregulated (heavy metal translocating P-type ATPase, CDF family zinc efflux transporter CzrB, Zn(II)-responsive metalloregulatory transcriptional repressor CzrA, cadmium-translocating P-type ATPase CadA) indicating a metal efflux strategy cells was being employed to detoxify their environment.

Taken together, these results indicate that an acetic acid environment causes cell stress and as a result, adapts by switching on numerous detoxification strategies to prevent oxidative membrane and DNA damage. The addition of Co-CQDs further stressed the cell with genes responding to oxidative stress being induced.

### Co-CQD nanoparticles cause multiple stresses on MRSA cells in an acetic acid environment

Buildup of cellular reactive oxygen species are closely linked to lipid peroxidation and cell wall damage consistent with our ultrastructural TEM and RNA sequencing conclusions. Given genes relating to combatting oxidative stress, in particular, peroxides were upregulated in Co-CQD NP treated samples in acetic acid conditions (i.e. protein arginine kinase (mscB), alkyl hydroperoxide reductase (ahpC) (**Figure 5E**) we next set out to phenotypically verify the transcriptional data. The oxidation of the antioxidant, ascorbic acid, in the presence of Co-CQD was initially measured. At concentrations between 12.5 – 100 µg/ml the addition of Co-CQD nanoparticle resulted in the dose-dependent oxidation of ascorbic acid by its reaction with ROS or autoxidation by cobalt (**Figure 5F**). Consistently, when treated bacteria were subjected to an ‘Amplex Red’ assay which measures hydrogen peroxide generation through the oxidation of 10-acetyl-3,7-dihydroxypenoxazine by peroxides in the presence of a horseradish peroxidase catalyst. The formation of increasingly red, fluorescent oxidation product was apparent with a stepwise increase with increasing nanoparticle concentration between 2.3 – 300 µg/ml verifying the production of peroxides (**Figure 5G**). In addition to peroxide generation, the particles were shown to produce superoxide anions (O2) through the fluorescent products of DHE oxidation by superoxide (**Figure 5H**). Acetic acid conditions increased the concentration of superoxide formed and superoxide dismutase, responsible for converting superoxide into H_2_O_2_ and O_2_, was upregulated in acidic conditions (2.2-Fold Change, 43.0 -Log (P)Adj). These experiments thus highlight that the Co-CQD particles potently induce multiple reactive oxygen species.

Alkaline phosphatase is produced and localized in the cell wall and cell membrane of bacteria. The detection of extracellular alkaline phosphatase is therefore a measure of cell wall damage after ROS damage. At both pH 7 and pH 5.5, there was an increase in extracellular alkaline phosphatase (AlkP) with increasing nanoparticle treatment with 0, 6, 30, 150, and 600 µg/ml showing AlkP dependent absorbance values of 0.00, 0.052, 0.055, 0.061, 0.092, and 0.164 respectively (**Figure 5I**). Bacterial cell wall damage was increased in acetic acid conditions (pH 5.5) with an additive effect from nanoparticle addition. At 600 µg/ ml treatment concentration, the extracellular alkaline phosphatase in the system was comparable to that of cells exposed to 15 mM H_2_O_2_ with both showing an absorbance of 0.164 and 0.166 respectively. At pH 7, alkaline phosphatase levels also increased with increasing nanoparticle concentration with 0, 6, 30, 150, and 600 µg/ml showing absorbance values of 0.000, 0.004, 0.011, 0.029, 0.049, and 0.166 respectively (**Figure 5I**).

Finally, bacterial membrane potential properties were determined using the cationic DiSC_3_(5) dye which quenches when it translocates and binds to the lipid bilayer of hyperpolarized membranes. Hyperpolarized membranes are increasingly negative in charge and hamper ion flux and intracellular signalling processes. This negative cellular surface charge may be due to carboxylic acid functional groups on Co-CQD interacting with the cell. At pH 7 increased Co-CQD treatment resulted in cells possessing increasingly negative membrane potential (hyperpolarization) resulting in stepwise fluorescence quenching of the DiSC_3_(5) dye compared to the untreated controls (**Supplementary Figure 5B**). At pH 5.5, this followed a logarithmic concentration dependent trend, with a trendline showing R^2^ values of 0.93 (**Figure 5J**).

Overall, Co-CQD nanoparticles cross MRSA cells membrane due to their relatively neutral charge, osmotic imbalance created from the presence of acetic acid which draws in water (and Co-CQDs) resulting in cell swelling. In the acidic environment, the particles generate ROS, particularly peroxide in nature but also superoxide and attack the cell from inside and out.

### Co-CQDs are biocompatible *in vitro* and *in vivo*

Having demonstrated the bactericidal effects of Co-CQDs on bacterial strains we then asked how Co-CQD affects mammalian cells and physiology. When Co-CQDs were added to wounds (1.09 mg/kg), there was no significant difference in mouse body weights over 7 days supporting the particles biocompatibility (**Figure 6A**). Further, representative images of wound closure over time are shown in **Figure 6B** demonstrated that addition of the Co-CQD had no visible toxicity on the wound healing process (**Figure 6B, C**). Combined this suggests that the Co-CQD were well tolerated by the mice. We next sought to assess their influence at a cellular level. During wound healing, fibroblasts generate collagen for promoting connective tissue formation^38^. The biocompatibility of the Co-CQD particles was assessed in human dermal fibroblasts at pH 7.0 and 5.5 at concentrations of 10, 50, 100, and 500 µg/ml. Human fibroblasts were resilient to Co-CQD treatments in both a neutral and acidic environment. At pH 7.0, dermal fibroblast cells had >96% viability between 0-500 µg/ ml, however, at 500 µg/ ml, the cells were smaller in size. At pH 5.5, Co-CQD treatments up to 100 µg/ml showed >95% viability, whilst at 500 µg/ml, the viability dropped to 76%. It is worth noting that at pH 5.5, cells did not proliferate but they did not die either. Once the acidic media (with and without Co-CQDs), were replaced with fresh unamended growth media, the cells were able to recover and proliferate (**Figure 6G**). This indicates that mammalian cells entered a quiescent state awaiting favourable conditions for proliferation with recovery growth rates indicating that their proliferative potential was not compromised. It is worth noting that lower molecular weight organic acids such as acetic acid (1%) has been shown to have a short lifetime in wounds (1 hour), before the pH returns to pre-treatment levels^20^. Therefore, the extreme 24 h exposure time in our experiments is unlikely to be experienced in *in vivo* conditions. An MBC of 38 µg/ml Co-CQDs was determined for MRSA in an aqueous acetic acid environment, but dermal fibroblasts were much more resilient showing 76% viability at >13-fold increased nanoparticle concentrations (500 µg/ml). Considering the high tolerance of dermal fibroblasts to Co-CQD treatment *in vitro*, we sought to assess if this also applied to *in vivo* models. Infected mice wounds treated with Co-CQDs were allowed to heal for 7 days and activated fibroblast derived fibrosis was measured through collagen 1 deposition in treated wound areas. In MRSA infected wounds, there was a significantly reduced collagen 1 deposition compared to the PBS control and Co-CQD treated MRSA infected wounds (**Figure 5 H, I**). This supports findings that MRSA wound infection impairs fibroblast function^39^. The Co-CQD treated wounds showed similar fibrosis content to that of the PBS control (**Figure 6H, I**), indicating that the treatment did not impair normal fibroblast function and collagen production.

**Figure 6:**
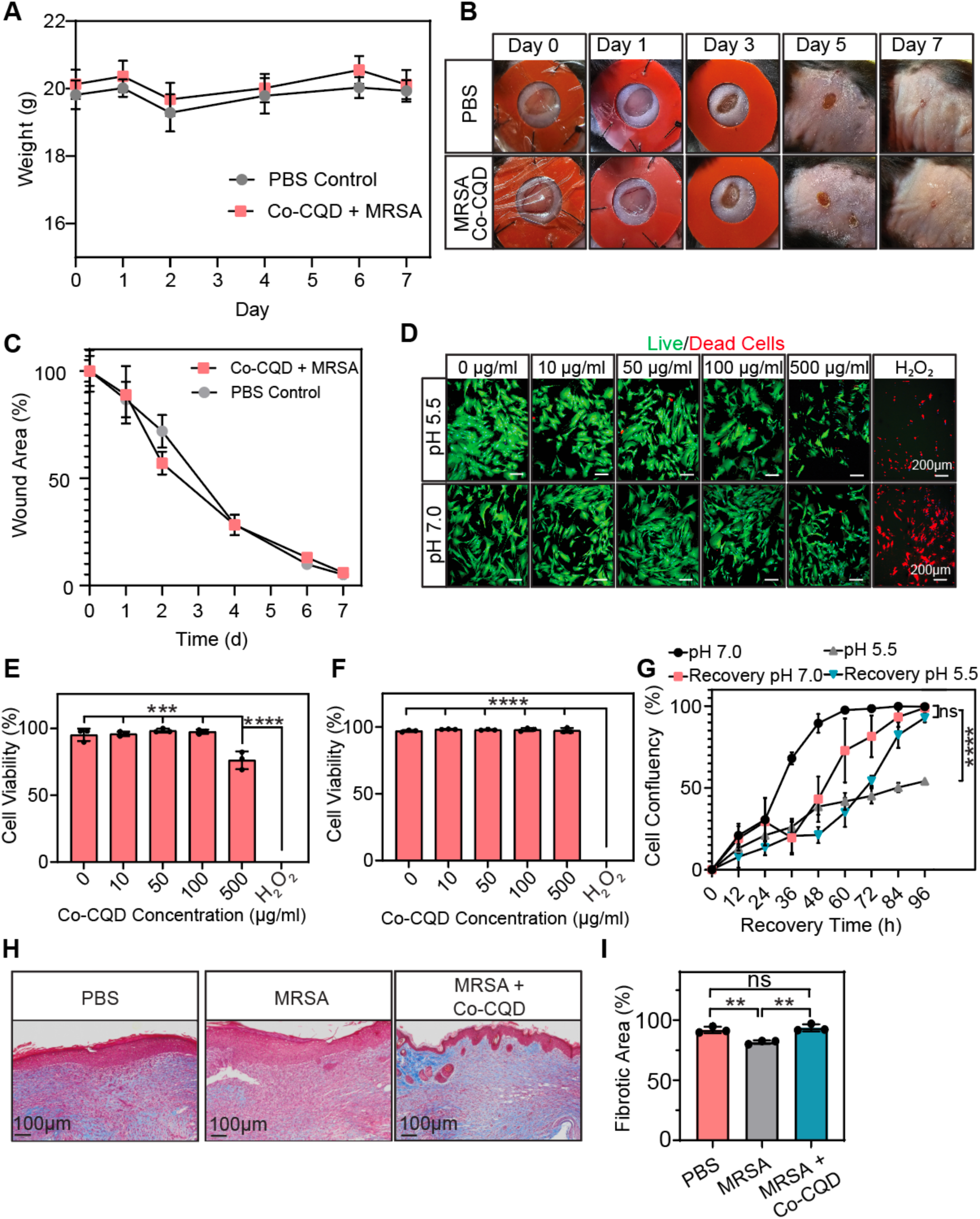
Co-CQD biocompatibility. **a**, Mouse weights over 7 days after inflicting full thickness wounds (5 mm diameter) in black 6 mice with and without Co-CQD treatment. **b**, Representative images of full thickness wounds in black 6 mice, treated with PBS or infected with MRSA and then treated with Co-CQDs for 7 d. **c**, quantification of wound closure following PBS and Co-CQD treated MRSA wounds over 7 d time period. **d**, Representative fluorescent confocal microscope images of human dermal fibroblasts in the presence of differing Co-CQDs concentrations at pH 7.0 and 5.5. The live/dead probes fluoresce green in viable cells and red in dead cells **e**, graphical quantification of aforementioned cell viability at pH 5.5 and **f,** at pH 7.0 when subjected to Co-CQDs at different concentrations over 24 h respectively. **G**, Cell proliferation (confluency over time) of human dermal fibroblasts subjected to pH 7.0, pH 7 + Co-CQDs (500 µg/ml), pH 5.5, and pH 5.5 + Co-CQDs (500 µg/ml) for 24 h followed by their recovery when growth media was replaced with unamended media following 24 h acetic acidic exposure (pH 5.5). Error bars represent the standard deviation. Statistical significance was determined using (b, f) a one-way analysis of variance (ANOVA) with Tukey’s multiple comparison test or a (d) Unpaired t test with Welch correction. ns, *, **, ***, **** signifies not significant, p < 0.05, p < 0.005, p < 0.0005 and p < 0.0001, respectively.

Combined, the addition of Co-CQD to wounds *in vivo* and dermal fibroblast *in vitro* demonstrated negligible toxicity. As such, Co-CQD are a viable and effective MSRA infection control strategy.

## Conclusions

In the present study biocompatible carbon quantum dot nanoparticles doped with cobalt were designed as potent antimicrobial agents against several bacterial pathogens including multi-drug-resistant gram-positive MRSA, gram negative *E. coli*, and vancomycin sensitive gram-positive *E. faecalis*. The particles exhibited strong antimicrobial activity against planktonic and surface colonised bacteria. In addition, the study demonstrates the impact of acetic acid on bacterial cell fitness and show that when Co-CQDs and acetic acid are used together, a synergistic, and greatly enhanced antimicrobial activity is achieved. Healthy healing wounds exhibit a fluctuating pH, but pathogen infections typically create a persistent alkaline environment slowing the healing process. Wound acidification has therefore been considered as a potential treatment for alleviating wound infection. In addition, an acidic environment has been shown to result in higher wound oxygen content, increased angiogenesis, and can transform bacterial end-products to less toxic forms (i.e. ammonia), promoting wound healing. In an acetic acid environment, MRSA and *E. coli* cells swelled and their osmotic balance was disrupted increasing the speed at which Co-CQDs could enter the cell. These particles generated peroxidase and superoxide ROS causing MRSA and *E. faecalis* to burst and *E. coli* to have their cell wall detach from the cytoplasm. Pathways analysis shows that Co-CQDs supressed bacterial protein translation and increased stress responses, mainly through oxidative damage. These findings were reinforced through cell swelling observations, extracellular alkaline phosphatase presence, capacity to generate peroxidase/peroxide and ascorbic acid oxidation by ROS. The particles were shown to be biocompatible with human dermal fibroblasts at concentrations >13 fold the MBC of MRSA in an acetic acid environment. When applied to mice, the particles removed MRSA infection and allowed wounds to heal at the same rate as uninfected wounds. Overall, this biocompatible Co-CQDs antimicrobial platform destroys bacteria through cellular destabilisation brought on by a weak acetic acid environment coupled with a reactive cellular destruction mechanism from the cobalt doped CQD nanoparticles.

## Methods

Cell culture and materials

Materials and reagents

### Synthesis of HACC CQD’s

Citric acid (5.4 g) and hexamine cobalt chloride (1.1 g) were dissolved in 30 ml of Milli-Q water and the solution was transferred to a poly(tetrafluoroethylene)-lined autoclave (50 ml). Vessels were heated at 205°C for 2 h and cooled to room temperature. The resulting red solution was purified via dialysis (100-500 MWCO) against ultrapure water (18,2 MΩ.cm resistivity) for 50 h with regular water changes. The purified particle solution was subjected to lyophilisation to obtain a fluffy, light purple powdered product.

### Characterization of CQDs

TEM images were obtained using a Hitachi HT7800 between 80-100 kV. Ultraviolet–vis absorption and fluorescence spectra were recorded using a Tecan SPARK® using 96 welled plates. For absorbance readings below 300 nm, a quartz cuvette was used in conjunction with an Agilent 8453 UV-visible spectroscopy system. FT-IR spectroscopy was measured using a Nicolet Protege 460 spectrograph between 400-4000 cm^−1^. Zeta potentials were measured using a Malvern 2000 Zetasizer within a DTS 1060C cuvette (Malvern) in Milli-Q water. To determine the nanoparticles cobalt concentration (Wt %), particles were digested in 2:1 nitric acid (70%): Sulphuric acid (99%) solution at 100°C for 10 h, followed by the slow addition of H_2_O_2_ (0.5 ml of 35% concentration) followed by 4 h further digestion. Particles cobalt concentration was analysed with HR-ICP-MS ElementXR, Thermo Fisher Scientific. X-ray photoelectron spectroscopy was run on a Thermo Scientific K-Alpha XPS with a monochromated Al 1487 eV Kα source. Peak fitting and background subtraction was interpreted using a Shirley background model using CasaXPS 2.3.23 software. Raman spectra was acquired on a Horiba Jobin Yvon LabRAM-HR800 system.

### Microbiological components

#### Strains and media

Methicillin and oxacillin resistant Staphylococcus aureus (MRSA, ATCC 43300), *Escherichia coli* (ATCC 25922), and vancomycin-sensitive *Enterococcus. Faecalis* (ATCC 29212) were routinely cultured in nutrient broth (NB)/ agar (NA) or brain/heart infusion (BHI) media.

#### Antimicrobial Assays (Liquid culture) (pH dependence) Metabolic activity (resazurin) & MIC

Overnight cultures of bacteria were reinoculated into fresh nutrient broth (1%) and grown to exponential phase. Nutrient broth (pH 5.5, 7.0, and 8.5) with differing Co-CQD treatment concentrations (300 – 9 µg/ml) were inoculated with exponentially growing bacteria (*E. coli* (2.65 × 10^6 CFU/ml), *S. aureus* 1.23 × 10^6 CFU/ml, *E. faecalis* 1.35 × 10^6 CFU/ml, and grown for 24 h at 37°C. Bacterial growth was determined by measuring absorbance at OD_600_). To determine the MBC, 10 µl of each growth culture was removed following the 24 h period, spread onto NA and incubated at 37°C. Following 48 h of growth, plates were assessed for colony growth.

To identify the metabolic activity of the growth culture, 30 µl of Resazurin solution (0.015% w/v) was added to 100ul of each 24 h exposed treatment. Following a 3 h incubation, the fluorescence of the solution was measured at ex 560 nm and em 590 nm. For samples with a pH of 5.5, the pH was adjusted by adding additional growth media (pH 7) prior to adding resazurin. This is because the acidic pH discolours the resazurin leading to inaccurate results.

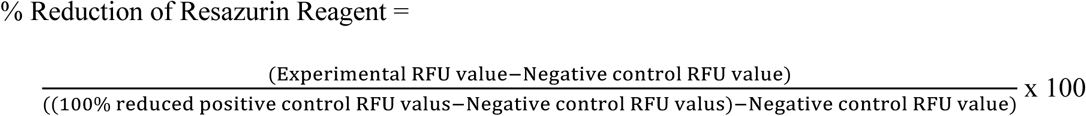

### Antimicrobial Assays (Solid growth)

#### Growth on plates with different initial bacterial seeding concentrations

Overnight cultures of bacteria were reinoculated into fresh nutrient broth (1%) and grown to exponential growth phase. A 10 µl aliquot of bacterial culture (E. coli (6.65 × 10^6 CFU/ml), S. aureus 5.11 × 10^6 CFU/ml, E. faecalis 5.46 × 10^6 CFU/ml was added to nutrient agar plates (pH 7.0 & 5.5) containing Co-CQD nanoparticles (300 −9 µg/ ml) in three independent replicates. Serial dilutions of the bacteria were added to the agar plate in the same manner to generate concentration dependant viability assay (10^6, 10^5, 10^4, 10^3 concentrations).

### TEM for HACC CD’s on bacterial cells

Samples were fixed in 2,5% glutaraldehyde (diluted in a 0,1M sodium cacodylate buffer) and 2% paraformaldehyde for 24 hrs at 4°C. Post fixation was performed for 1h (on ice) in 1% osmium tetroxide (EMS # 19134) diluted in 0,1 M sodium cacodylate buffer, followed by 2 washing steps. The samples were then dehydrated using a graded ethanol series (30%, 50%, 70%, 96% and 100%) before transferred to a 1:1 solution of 100% ethanol:propylene oxide (15 min). Samples were then transferred to 100% propylene oxide (15min) before gradually introducing agar 100 resin (AgarScientific R1031) drop by drop over the next hours. Samples were then transferred to a small drop of 100% resin and excess propylene oxide was allowed to evaporate (1h). Samples were then transferred to 100% resin and placed in molds and left in room temperature overnight. The molds were placed at 60℃ for 48h to polymerize. Ultra-sections of approximately 60nm were placed on 100 mesh formvar coated (EMD # 15820) copper grids (EMS #G100H-Cu) and stained with 2% uranyl acetate (EMS # 22400) and lead citrate (VWR #1.07398). Grids were imaged using a Hitachi HT7800 transmission electron microscope at 100kV.

### Bacterial cell size

The width and height of all in-tact bacteria in imaged under TEM (at least 3 images from independent samples per strain, treatment and pH) were manually measured using measurement tools from the Hitachi EMP-EX software. Peptidoglycan thickness from gram positive species was also measured.

### Cell-cell interactions (*E. coli*)

From TEM images, each *E. coli* cell was recorded as either touching another cell or not. The number of cells that were touching at least one other cell was represented as a percentage of total cells per TEM image (at least 3 images from independent samples for E. coli per treatment and pH).

### RNA-seq data analysis

Methicillin and oxacillin resistant Staphylococcus aureus were grown in unamended nutrient broth (NB) to an absorbance of OD_600_ = 2.5. This solution was transferred into sterile 1ml tubes, centrifuged and the supernatant was removed. The bacteria were centrifuged, and the supernatant was removed. The bacterial pellet was resuspended with 1 ml of the following solutions and incubated at 37°C for 3 h:

1. NB unamended
2. NB pH 5.5 (acetic acid adjusted)
3. NB pH 5.5 (acetic acid adjusted) + Co-CQD (600 µg/ml)

RNA was extracted using TRIzol™ Max™ Reagent and Phasemaker™ Tubes Complete System (A33251) as per the manufacturer’s instructions (Pub. No. MAN0016163, Rev. A.0). Crude RNA was purified twice using RNA Clean & Concentrator-25 kit (Zymol). Ribosomal RNA removal, cDNA library construction, and paired-end sequencing were performed using the NextSeq 500 platform (150 cycles (2 x 75 bp paired end reads), ca. 100 M read pairs by NorSeq Sequencing core (Ullevål) (Oslo University Hospital, Norway).

Sequenced reads were quality checked with FastQC and aligned to the Refseq reference genome (GCF_003052445.1) using Bowtie2. Aligned reads were counted and summarized for the annotated genes using featureCounts. Differential gene expression analysis was performed by DESeq2. Differential expressed protein-coding genes with a log2 fold-change > 0.6 or < −0.6 and adjusted p-value < 0.05 were used for pathway analysis using DAVID^40^.

### STRING analysis

To investigate the protein-protein functional interactions amongst genes identified in the pathways analysis, STRING 12.0 database was used. Proteins from each significant pathway were entered into the database and *S. aureus* was chosen as the species of reference.

### Alkaline phosphatase

S. aureus was grown to an OD_600_ of 0.2 (5 ml) and HACC CD was added within the range of 1000 – 0.2 µg/ml at pH 5.5 and 7.0 (15mM H_2_O_2_ serving as the positive killing control) for 6 h. The solution was centrifuged at 10,000 X G for 10 mins and 1 ml of the supernatant was mixed with 2-amino 2-methyl 1-propanol buffer (pH 10.5) containing paranitrophenyl phosphate (PNPP) at 2 mg/ml concentration. The samples were incubated at 30°C for 30 mins and absorbance was measured using a quartz cuvette.

### ROS generation with ascorbic acid (AA)

Ascorbic acid can be oxidised by ROS forming dehydroascorbic acid. Co-CQDs capacity to generate ROS at different pH’s was determined by measuring a decrease in absorbance from ascorbic acid (absorption peak at 266 nm) when in the presence of Co-CQQDs. PBS buffer was adjusted to pH 5.5 and 7.0 with ascorbic acid and Co-CQDs were introduced at concentrations ranging from 12.5 – 100 µg/ml. The solutions were incubated for 2 h at 37°C and ascorbic acids characteristic peak at OD_266_ was measured in a quartz cuvette. Above 100 µg/ml Co-CQD concentrations, the AA absorbance peak at OD_266_ was overshadowed by the absorbance of Co-CQD particles and so, only low HACC CD concentrations were measured.

### Superoxide with Nucleic Acid

To investigate the capacity of Co-CQDs to generate superoxide radicals, dihydroethidium (DHE)was used as the fluorescent probe. Dihydroethidium was dissolved in DMSO and then resuspended in PBS to make a working solution of 10 µM. MRSA cells (1 ml, OD_600_= 0.2) were subjected to Co-CQDs 1.2 – 150 µg/ml at 37°C for 1h. The cells were centrifuged, and the pellet was retained and incubated in DHE working solution for 30 mins. The cells were then centrifuged, washed three times with PBS and fluorescence was measured at λex/λem = 518 nm/606 nm.

### Membrane polarisation

MRSA, *E. coli*, and *E. faecalis* (25 µl, OD_600_ = 0.10) were added to 125 µl of 3,3′-dipropylthiadicarbocyanine iodide (diSC_3_-5) made to a concentration of 1.0 µM in HEPES buffer (10 mM) s supplemented with glucose, KCl and MgSO_4_ at pH 5.5 and 7.0. The solution was allowed to equilibrate for 30 m before the addition of Co-CQDs (37.5 – 1000 µg/ml). The fluorescence was monitored at 2 h using a microplate reader (Tecan Spark), λex/λem = 622 nm/674 nm.

### Hydrogen Peroxide generation assay (Amplex® Red)

To determine the generation of hydrogen peroxide from Co-CQD nanoparticles, an Amplex Red assay was conducted as per manufacturer’s instructions (Amplex Red Kit: Invitrogen A22188). Briefly, Co-CQD nanoparticles (2.3 – 300 µg/ ml) were incubated with 100 µM Amplex® Red reagent and 0.2 U/mL horseradish peroxidase for 30 minutes in the dark at room temperature. The positive control consisted of 15 mM H_2_O_2,_ and the negative control was pure water. Fluorescence spectra indicative of H_2_O_2_ production was measured at λex/λem = 560 nm/580-650 nm.

### Live/Dead

Human Dermal fibroblasts (dFIB) (CC-2509- LONZA) were cultured in DMEM (10% FBS and 3% glutamine. Cells were grown to between 50-90% confluency in IBIDI 8 welled plates and Co-CQD nanoparticles were added at different concentrations (0, 10, 50, 100 & 500 µg/ ml) and incubated for 24 h at pH 7 and 5.5 (acetic acid adjusted). Hydrogen peroxide (15 mM) was the positive killing control. Cytotoxicity was determined by a LIVE/DEAD® Cell Imaging assay (R37601-Invitrogen). Briefly, the kit reagents were added to cells as per manufacturers suggestions, incubated for 20 minutes and imaged on a Dragonfly 505 (Andor Technologies, Inc) using 60 X objective with oil immersion. Images were taken using excitation/ emission wavelengths 488/515 and 570/ 600 nm.

### Fibroblast Proliferation

Human Dermal fibroblasts (dFIB) (CC-2509- LONZA) cultured in DMEM (10% FBS and 3% glutamine) were grown to 80-90% confluency in Incucyte® ImageLock 96-well plates (Sartorius) and 700-800 µm scratch was made in cells using an Incucyte® 96-Well Woundmaker. The cells were washed with PBS and then exposed to growth media at pH 7.0 and 5.5 (acetic acid adjusted) containing Co-CQD nanoparticles at 500 µg/ ml for 24 h. Following this, the media was removed and replaced with fresh, unamended media for a further 72 h. The cells were incubated and imaged in an IncuCyte® Live-Cell Analysis System. Confluence was measured by the area of the scratch filled with migrating cells over time.

### Antimicrobial evaluation in mice

Mouse experiments were approved by the Norwegian Food and Animal Safety Authority (Mattilsynet) FOTS ethics approval # 30052. All surgeries were performed under general anaesthetic conditions to minimise suffering and mice were randomly assigned for analysis.

The dorsal of Female C57BL/6 mice (8-10 weeks) were shaved and hair was further removed using Veet hair removal cream. A single punch biopsy wound was made on each mouse (5 mm). The wounds were infected with MRSA (1.3 × 10^^7^ CFU) and allowed to dry for 3 h followed by the addition of Co-CQD nanoparticles in PBS at pH 5.5 (0.87 mg/kg +/− 10%). The wounds were kept from contracting by gluing a splint around the wound. The wound was then covered by a commercial transparent wound dressing (Tegaderm Film, 3M). The mice (each treatment n=6) were sacrificed after 24 h and their wounds were removed with scissors. Three wounds were homogenised and mixed in 1 ml sterile PBS, and 10 µl of this was spread onto agar plates. The remaining 3 wounds were collected, and paraffin embedded for H&E staining and antibody labelling.

### Wound healing evaluation in mice

The dorsal of Female C57BL/6 mice (8-10 weeks) were shaved and hair was further removed using Veet hair removal cream. A single excisional skin wound was made as previously described. The wounds were infected with MRSA (2.2 × 10^^6^ CFU) and allowed to dry for 3 h followed by the addition of Co-CQD nanoparticles in PBS at pH 5.5 (1.09 mg/kg +/− 10%). The wounds were kept from contracting by gluing and suturing a silicone splint around the wound. The wound was then covered by a commercial transparent wound covering (Tegaderm Film, 3M). The wound covering was removed following 24 h and the splint was removed after 3 days (following scab formation). Mice were weighed every 1-2 days for 7 days and wound lengths and widths were measured with electronic callipers. The mice (each treatment n=6) were sacrificed after 7 days, and their wounds were removed with scissors. Three wounds were homogenised and mixed in 10 ml sterile PBS, and 10 µl of this was spread onto agar plates and incubated at 37°C for 48 h. The remaining 3 wounds were collected, and paraffin embedded for H&E staining and antibody labelling.

### H&E Staining

Samples were fixed in formalin (10%), dehydrated through increasing ethanol steps (70-100%) and then cleared in xylene before paraffin infiltration. Samples were then embedded and sectioned. The HE (hematoxylin and Eosin) staining included removing the paraffin (dewaxing), rehydration, staining with Hematoxylin, rinsing, staining with Eosin, rinsing, dehydration, clearing and mounting. Samples were then imaged using the Olympus VS120 slide scanner.

### Tissue Immunofluorescence

For tissue immunofluorescence, 5 mM tissue sections were deparaffinized in xylene (10 mins twice) and rehydrated by immersing in a series of graded alcohols (100% EtOH 10 mins twice, 95% EtOH for 10 mins once, 70% EtOH for 10 mins twice) and distilled water (5 mins twice). Antigen retrieval was performed by boiling the sections in a pressure cocker at 98°C for 12 min. For staining of MPO and iNOS, antigen retrieval was performed using Tris-based antigen unmasking solution, pH 9.0 (Vector Laboratories, H-3301) whereas antigen unmasking for F4/80 was performed in sodium citrate buffer, pH 6.0 (Vector Laboratories, H-3300). After cooling at RT for 40 mins, sections for F4/80 were blocked in 10% normal goat serum (Invitrogen, 50197Z) whereas 5% Bovine serum albumin (BSA) diluted in PBS was used for MPO and iNOS. Following blocking for 1 hour at room temperature, slides were incubated at 4°C overnight with primary antibodies. F4/80 (Cell Signalling, 70076) and MPO (R&D systems, AF3667) were diluted 1:200, whereas iNOS (Abcam, ab15323) was diluted 1:100, respectively. After washing in PBS containing 0.05% Tween-20, samples were incubated for 1 hour in the dark with secondary antibody, diluted 1:200 (Alexa-Fluor 647: Invitrogen, A21244, Alexa-Fluor 546: Invitrogen, A11056). After final washing, sections were mounted using Prolong Diamond antifade reagent containing DAPI (Invitrogen, P36970). Images were acquired using an Olympus VS120 slide scanning system (40x objective).

### Trichrome stain

Collagenous connective tissue fibres were visualized using the Trichrome Stain kit from Abcam (ab150686). In brief, 5 mM sections were deparaffinized and rehydrated as described under the tissue immunofluorescent section. Next, slides were stained according to the manufacturer’s instructions. Slides were cleared in xylene and mounted with Eukitt Quick-hardening mounting media (Sigma-Aldrich, 03989). Images were obtained on the Olympus VS120 slide scanning system (40x objective).

### Statistics and reproducibility

All microscopy including TEM and confocal images were independently repeated in triplicate. Error bars are representative of standard deviation from the mean. Mice were randomly assigned to treatment groups and randomly assigned to wound bacterial load or wound histology investigations. Statistical analysis between treatment groups were determined by one way or two-way analysis of variances, complimented by Tukey’s post hoc analysis. In certain assays, unpaired t test with Welch correction were employed. All statistical analysis used a 95% confidence interval (p-value < 0.05).

## Acknowledgements

We would like to acknowledge the Molecular Imaging Centre (MIC), Department of Biomedicine, University of Bergen for use of their transmission electron microscope and confocal microscopes. In addition, we would like to thank the EarthLab ICP-Laboratory, Department of Earth Science, University of Bergen for ICP-MS analysis. We thank Madeleine Kersting, QIMR Berghofer Medical Research Institute for help with illustrations. Additional project financial support was provided by the Familien Blix Fond (Project#: 103267101) and The General Medical Research Fund (Project #: 103520115).

