## Supplemental Figures for "Acetic acid induces osmotic imbalances in drug-resistant bacteria synergistically enhancing cobalt-doped carbon quantum dots bactericidal efficiency"

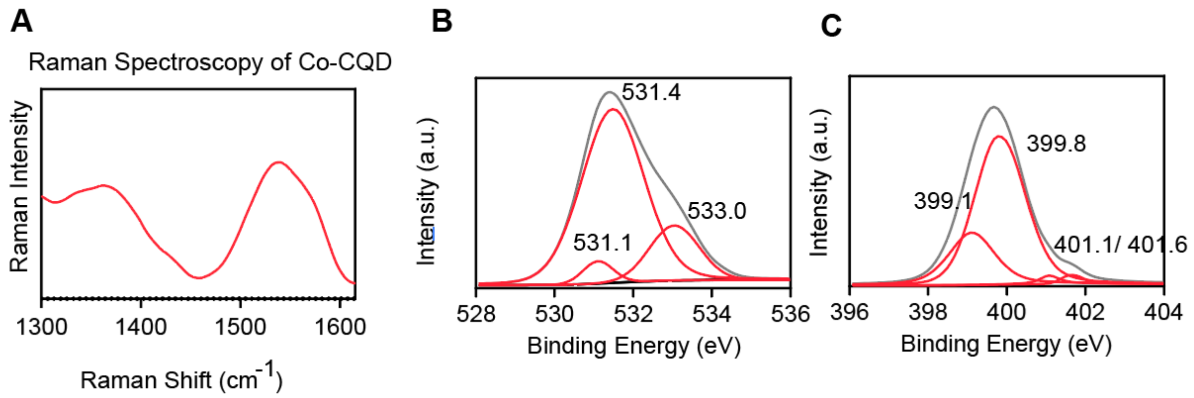


**Supplementary Figure 2: Co-CQD characterisation: a**, Raman spectra of Co-CQDs between 1300-1600 $cm^{-1}$. **b**, X-ray photoelectron spectroscopy (XPS) of O1s and **c,** N1s peaks from Co-CQDs.

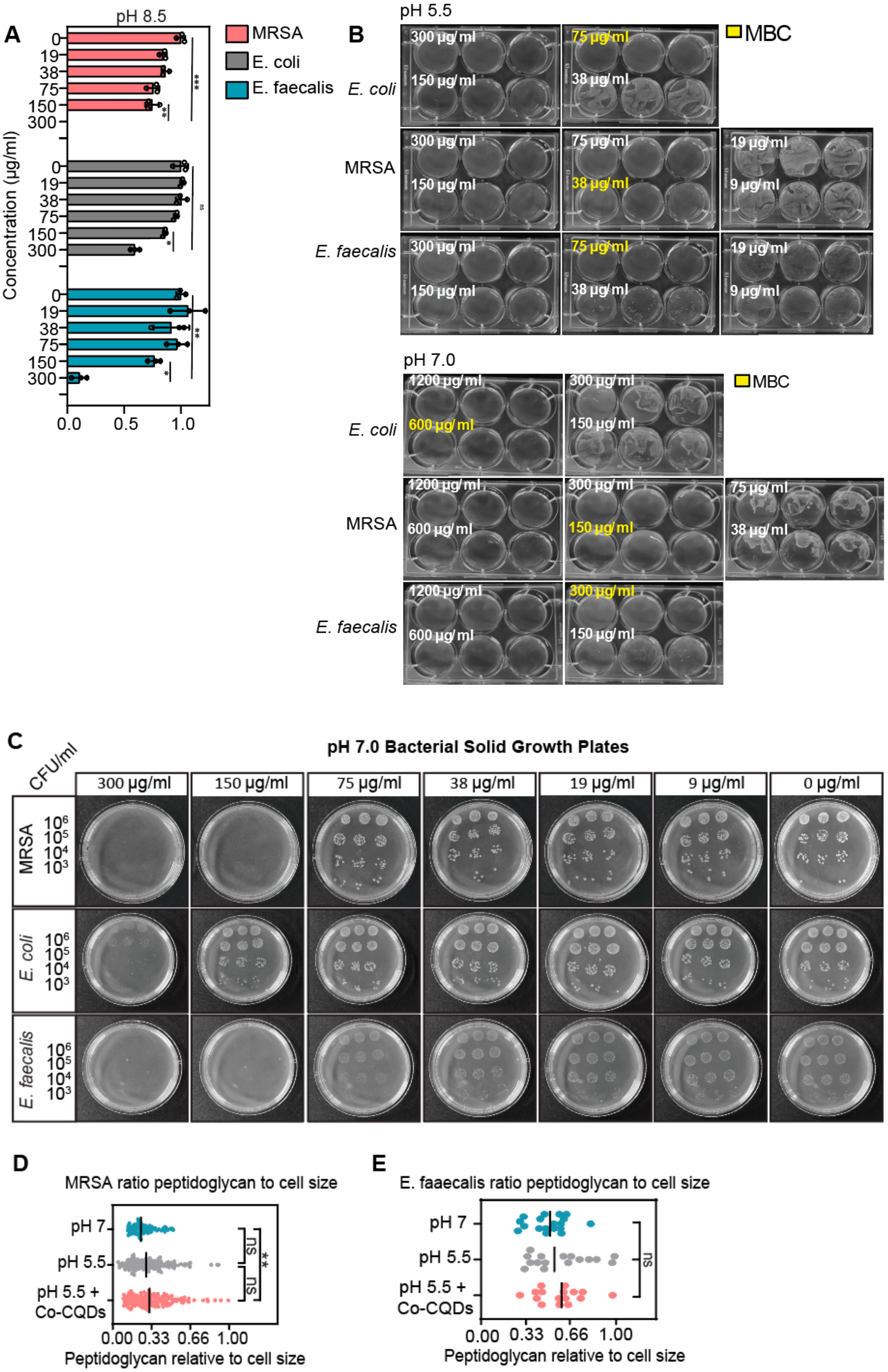
A
pH 8.5
MRSA
E. coli
E. faecalis
Concentration (μg/ml)
0
19
38
75
150
300
0.0
0.5
1.0
***
**
ns
*
B
pH 5.5
MBC
E. coli
MRSA
E. faecalis
300 μg/ml
150 μg/ml
75 μg/ml
38 μg/ml
19 μg/ml
9 μg/ml
pH 7.0
1200 μg/ml
600 μg/ml
C
CFU/ml
pH 7.0 Bacterial Solid Growth Plates
300 μg/ml
150 μg/ml
75 μg/ml
38 μg/ml
19 μg/ml
9 μg/ml
0 μg/ml
MRSA
E. coli
E. faecalis
10^6
10^5
10^4
10^3
D
MRSA ratio peptidoglycan to cell size
pH 7
pH 5.5
pH 5.5 +
Co-CQDs
ns
**
0.00
0.33
0.66
1.00
Peptidoglycan relative to cell size
E
E. faaecalis ratio peptidoglycan to cell size
pH 7
pH 5.5
pH 5.5 +
Co-CQDs
ns
0.00
0.33
0.66
1.00
Peptidoglycan relative to cell size

**Supplementary Figure 3: Antibacterial activity of Co-CQDs. a**, Optical density ($OD_{600\ nm}$) of MRSA, *E. coli*, and *E. faecalis* in pH adjusted liquid growth cultures after 24 h exposure to Co-CQDs at pH 8.5. **b**, MBC of MRSA, *E. coli*, and *E. faecalis* from the aqueous growth assay supplemented with differing Co-CQD concentrations at pH 5.5 (0 – 300 µg/ml) and pH 7.0 (0 – 1,200 µg/ml). **c**, Growth of MRSA, *E. coli*, and *E. faecalis* on nutrient agar plates supplemented with differing Co-CQD concentrations (0 – 300 µg/ml) at pH 7.0. Each plate has been seeded with different concentrations of bacteria (5.1 X $10^6$, $10^5$, $10^4$, and $10^3$ CFU/ ml) from top to bottom in triplicate, **d**, Ratio of *MRSA* peptidoglycan content relative to cell size **e**, Ratio of *E. faecalis* peptidoglycan content relative to cell size

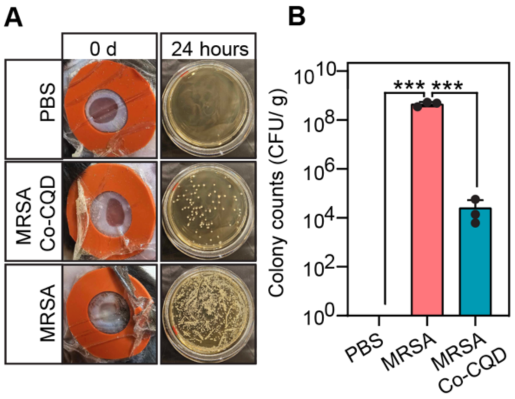


**Supplementary Figure 4: Co-CQD treatment removes infection in vivo. a**, Wound areas following 24 h incubation with PBS, Co-CQD + MRSA and MRSA with agar plates representing their wound bacterial load after 24 h exposure. **b**, quantification of bacterial presence in wound areas following treatments after 24h. Statistical significance was determined using (b, f) a one-way analysis of variance (ANOVA) with Tukey's multiple comparison test or a (d) Unpaired t test with Welch correction. ns, *, **, ***, **** signifies not significant, $p < 0.05$, $p < 0.005$, $p < 0.0005$ and $p < 0.0001$, respectively.

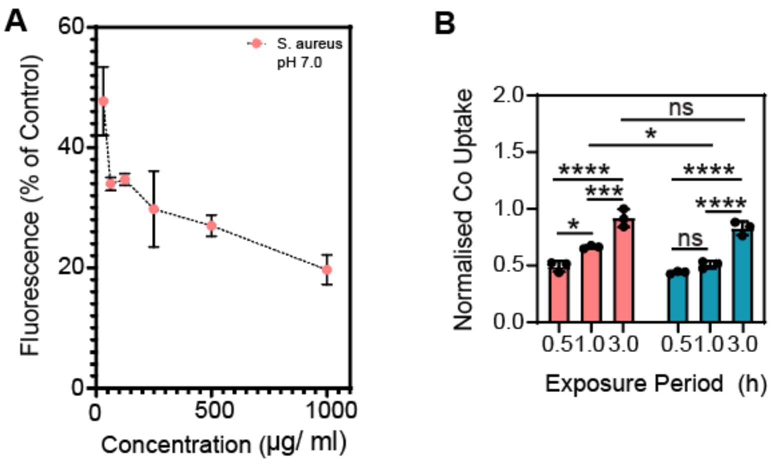


**Supplementary Figure 5: Mechanism of Co-CQDs Antibacterial activity. a**, Membrane hyperpolarisation of MRSA at pH 7.0 resulting from differing Co-CQD exposure. **b**, Normalised uptake of Co-CQD at pH 5.5 and 7.0 measured through cobalt concentration at 0.5, 1, and 3 h time periods. Statistical significance was determined using (b, f) a one-way analysis of variance (ANOVA) with Tukey's multiple comparison test or a (d) Unpaired t test with Welch correction. ns, *, **, ***, **** signifies not significant, $p < 0.05$, $p < 0.005$, $p < 0.0005$ and $p < 0.0001$, respectively.
